# Pan-viral protection against arboviruses by targeting inoculation site-based skin macrophages

**DOI:** 10.1101/566885

**Authors:** Steven R Bryden, Marieke Pingen, Daniella A Lefteri, Jack Major, Leen Delang, Sofie Jacobs, Rana Abdelnabi, Johan Neyts, Janne Miltenburg, Henna Khalid, Andrew Tuplin, Andres Merits, Emilie Pondeville, Julia Edgar, Gerard J Graham, Kave Shams, Clive S McKimmie

## Abstract

Arthropod-borne viruses (arboviruses) are important human pathogens for which there are no specific antiviral medicines. The large number of genetically-distinct arbovirus species, coupled with the unpredictable nature of their outbreaks, has made developing virus-specific anti-viral medicines challenging. Instead, we have defined and targeted a key aspect of the host innate immune response to virus at the arthropod bite that is common to all arbovirus infections, potentially circumventing the need for virus-specific therapies at this site. Using mouse models and human skin explants, we identify innate immune responses by dermal macrophages in the skin as a key determinant of disease severity. Post-exposure treatment of the inoculation site by a topical innate immune agonist significantly suppressed both the local and subsequent systemic course of infection and improved clinical outcome in mice to infection with a variety of arboviruses from the Alphavirus, Flavivirus and Orthobunyavirus genuses. In the absence of treatment, anti-viral interferon expression to virus in the skin was restricted to dermal dendritic cells. In contrast, targeting the more populous skin-resident macrophages with an immune agonist elicited protective responses in key cellular targets of virus that otherwise replicated virus to high levels. By defining and targeting a key aspect of the innate immune response to virus at the mosquito bite site, we have shown that it is possible to improve outcome to infection by targeting pathways activated at the site of inoculation, and thereby identified a putative new strategy for limiting disease following infection with a variety of genetically-distinct arboviruses.

**One-sentence summary:** We demonstrate that activation of innate immune responses to arbovirus at the mosquito bite is a limiting factor for preventing efficient systemic dissemination of virus and that therapeutic targeting of skin-resident macrophages can have defining inhibitory effects on the later systemic course.

## Introduction

Emerging and re-emerging arboviruses pose an increasing threat to human health. There has been a substantial increase in both the incidence and geographical range of medically-important arboviruses spread by mosquitoes, which infect hundreds of millions of people each year and include the Zika (ZIKV), dengue (DENV) and chikungunya (CHIKV) viruses. Arboviruses are a large, genetically-diverse group of viruses that cause a wide spectrum of diseases in humans (1–5). Despite their genetic diversity, it is nonetheless difficult to clinically differentiate between these infections in the early stages of disease, as they are either asymptomatic or present as a non-specific febrile viral illness. In many geographic areas this is compounded by the widespread co-circulation of distinct species of arboviruses in the same geographic area (6). Together, these factors complicate the use of putative virus-specific anti-virals, which for acute infections are most often only efficacious when given during early stages (7). This, when combined with our inability to accurately predict the timing and location of future epidemics (8), makes stockpiling future virus-specific drugs and vaccines difficult. Currently, there are few vaccines and no antivirals available for arbovirus infections. We suggest that due to the diversity of arbovirus genetics, their common clinical features and their unpredictable epidemiology, the development of a pan-viral medicine that is efficacious for multiple arbovirus infections would be highly advantageous.

Infected mosquitoes deposit virus into the skin dermis as they probe for a bloodmeal, triggering activation of distinct inflammatory pathways in response to mosquito biting and to virus sensing (9–11). We and others have demonstrated that host responses in the skin to mosquito biting, or mosquito saliva, has a defining influence on the systemic course of infection for a wide-variety of genetically-distinct arboviruses including flaviviruses, alphaviruses and bunyaviruses (10, 12, 13). As such, this is a key stage of infection during which virus replicates rapidly before disseminating to the blood and remote tissues. However, some of the original virus inoculum also disseminates directly to the blood and the draining lymph node (dLN) where replication is also established prior to systemic dissemination (9, 14). Thus, it is not clear what role skin-specific innate immune responses, activated by virus sensing at the mosquito bite, has on modulating the subsequent systemic course. In this study, we wanted to define the relevance of skin virus-sensing pathways and determine whether this can be targeted to modulate outcome of infection.

Innate immune sensing of virus activates immune pathways that are distinct to those activated by mosquito biting, resulting in expression of type I interferons (IFN) and the anti-viral genes they upregulate, IFN-stimulated genes (ISG). Despite the evolution of multiple strategies by arboviruses to counteract IFN they are nonetheless highly sensitive to them, suggesting that therapies designed to target these pathways could be generalisable to a range of viruses (15–18). Systemic administration of type I IFN has been used to treat HCV, although febrile-like side effects are common (19) and as such it is not suitable for e.g. long-term prophylactic use. However, previous attempts to inhibit arbovirus infection by systemic administration of innate immune agonists or type I IFN have only been efficacious when given prior to infection (20–24). In contrast, we suggest targeting processes at the inoculation site, which represents a discreet identifiable locale that can be targeted post-mosquito bite, prior to the systemic dissemination of virus. However, targeting these pathways is challenging as the coordination of early innate immune responses to virus at mosquito bites are not well defined.

In this study, we define key aspects of host response to virus at mosquito bites and demonstrate that this can be therapeutically modulated to suppress viral replication and the development of clinical disease. We suggest that for virus to successfully disseminate from the skin to blood at a sufficient level to induce a high-titre viremia, sufficiently robust levels of replication must occur at the inoculation site before an anti-viral state that follows host IFN production. In the absence of treatment, successful dissemination of virus from the skin was associated with sub-optimal activation of local IFN responses. Using an immune-competent mouse model of arbovirus infection and *ex vivo* infection of human skin explants, we demonstrate that therapeutic manipulation of skin IFN pathways by post-exposure topical application of a widely-used generic innate immune agonist, protected against infection with multiple genetically-distinct arboviruses. In doing so, we have defined key aspects of skin innate immune response to both virus and innate immune agonists at the mosquito bite, including the identification of a population of skin-resident macrophages that are sufficient to mediate agonist-induced protection from infection. Together, this work demonstrates that early events of infection at the mosquito bite are pivotal for defining the subsequent systemic course and that this site can be targeted through post-exposure therapeutic intervention.

## Results

### Skin innate immune responses to virus infection at the inoculation site are a key determinant of the systemic course

Studying innate immune responses at the tissue- and system-wide level to arboviruses that are medically-important in humans, has been frequently complicated by their inability to replicate in immunocompetent mice (10). Therefore, we chose to primarily study host innate immune responses to a prototypic model arbovirus SFV (genus Alphavirus). Unlike most human arboviral pathogens, SFV is capable of efficiently replicating, disseminating systemically within, and causing clinically-observable disease in immunocompetent mice. SFV is a close relative of CHIKV (25) that has a large number of genetically-modified clones and has been used extensively to study host response to infection. SFV replicates quickly following infection of mouse skin, with viremia peaking by 24 hours post-infection (hpi). Dissemination of SFV to brain tissue can result in encephalitis, neurological signs and death (9, 26).

We wanted to determine whether targeting early innate immune responses in the skin had any observable effect on the later systemic course of infection. However, it was first necessary to define the kinetics and magnitude of endogenous host ISG responses to infection with SFV6, in the absence of therapies. Following infection of the foot skin, ISG expression in response to virus infection was slow and of low-magnitude compared to induction of ISG in the dLN, with ISG expression not significantly elevated until 24 hpi, with levels peaking at 48 hpi (Fig 1A-D). In comparison, the dLN rapidly and robustly upregulated ISG, with levels peaking by 16 hpi. Thus, upregulation of anti-viral ISG in the skin could be detected only after virus had disseminated systemically.

**Fig. 1.**
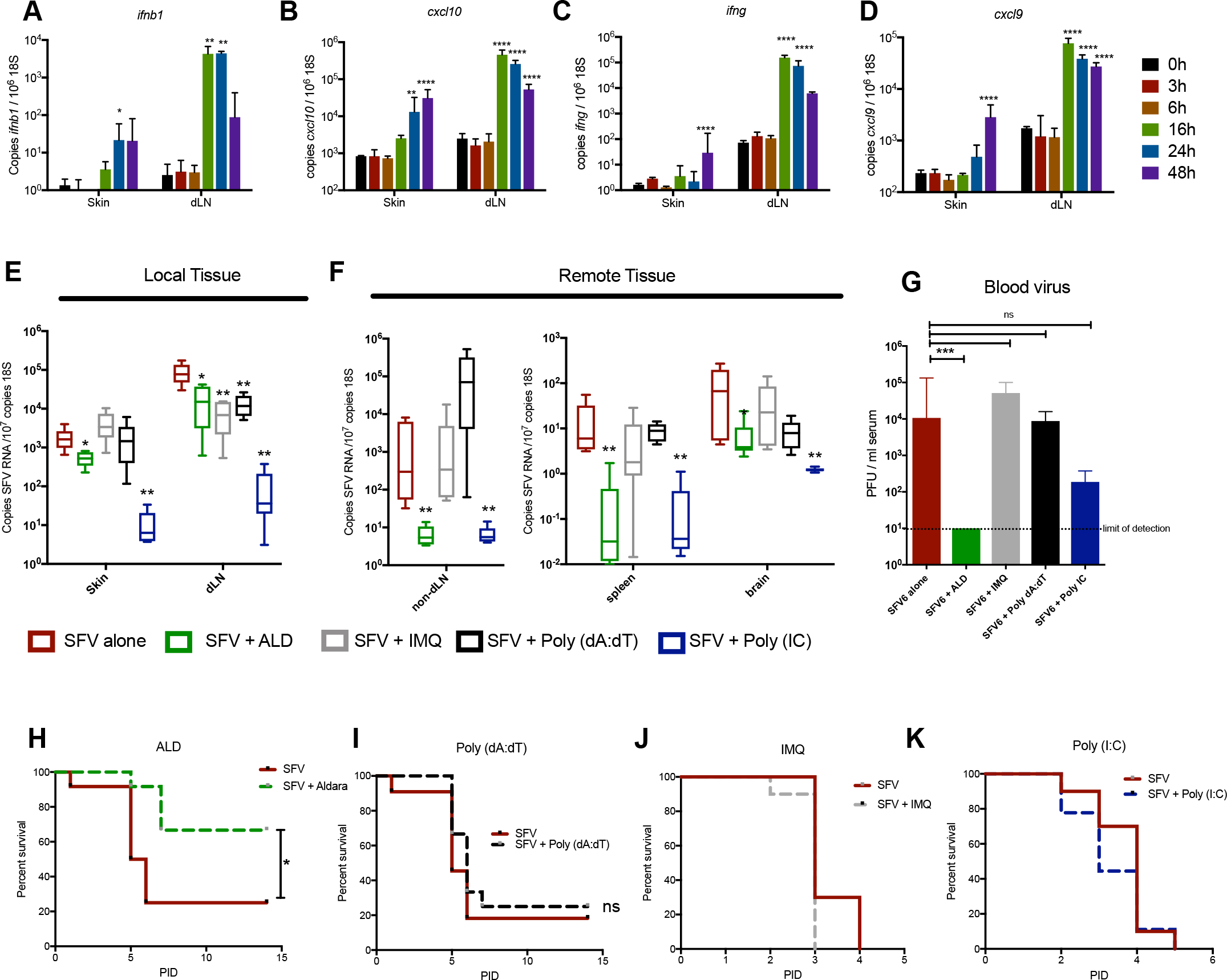
Immunomodulation of the skin inoculation site enhances host resistance to virus infection. (A-D) Mice were infected subcutaneously (s.c.) with 250 PFU SFV6 in the upper skin of the left foot. Copy number of host (A) *ifnb1*, (B) *cxcl10* (C) *ifng* (D) *cxcl9* and 18S RNA in skin and draining popliteal lymph node (dLN) was determined by qPCR (n=6). (E-G) 1h prior to infection with SFV6, mice were treated with either topical IMQ (Aldara, ALD) or injected s.c. with 6μg of either; aqueous IMQ, poly(dA:dT) or poly(I:C). Copy number of SFV RNA and host 18S was determined by qPCR at 24 hpi. Non-draining LN was the popliteal LN contralateral to infection. (G) Virus titers in the serum were quantified by plaque assay (n=6). (H-K) Mice were left resting or pre-treated for 1h with; (H) 2mg topical ALD (n=12); (I) s.c. 6μg Poly (dA:dT) (n=12); (J) s.c. 6μg IMQ (n=10); (K) s.c. 6μg Poly (IC) (n=10); and then infected with SFV6 viruses. PID=post infection day. Bars significantly different to 0 h or SFV alone respectively, *p<0.05, **p<0.01, ***p<0.001, ns=not significant (Kruskal-Wallis and logrank (Mantel Cox) test)

It is not clear what relevance skin-specific innate immune responses to arbovirus have on the subsequent systemic course of infection, nor whether virus replication in the skin is absolutely necessary to establish viremia. To determine whether therapeutic induction of anti-viral innate immune pathways at the inoculation site could have any effect on the systemic course of infection, we injected or topically applied a range of distinct innate immune agonists to a defined cutaneous site one hour prior to infection with SFV6 (Fig 1E-K). The TLR7 agonist imiquimod (IMQ) was administered either as a subcutaneous (s.c.) injection or applied topically as a cream (Aldara™, ALD). At 24 hpi, only topical ALD and poly(I:C)-treated mice exhibited lower viral RNA at the skin inoculation site, while all innate immune agonists decreased virus in the dLN (Fig 1E). Poly(I:C) was particularly potent at decreasing viral RNA in most tissues analyzed, although viremia (Fig 1G) and survival to infection (Fig 1K) was not significantly affected. In comparison, only ALD significantly; decreased viral replication at both local tissues (skin and dLN); tissues remote from the inoculation site (non-dLN, spleen and brain); reduced infectious virus in blood; and limited the development of clinical disease (Fig 1H). As SFV6 is highly virulent in mice, it does not model all arboviral disease seen in humans that are often non-fatal. Therefore, to better model the human situation we also tested the ability of IMQ to suppress infection with SFV4, a less virulent strain than SFV6 (Fig S1A). Again, topical application of ALD at the site of SFV4 inoculation significantly suppressed infection.

### Post exposure targeting of the inoculation site by topical application of an innate immune agonist protects mice from infection with virus

Future treatment modalities that target early stages of infection are likely to involve post-exposure treatment, e.g. once the erythema of a bite is apparent. Because pre-bite application of ALD was most efficacious in increasing host resistance to infection, we next determined its efficacy when applied post-infection. In addition, as arbovirus infection of skin always occurs in the context of an arthropod bite, we used a mouse model that additionally incorporates biting *Aedes aegypti* mosquitoes. Host response to mosquito bites include edema and an influx of leukocytes that enhances host susceptibility to infection with virus (9, 10, 27). In this model, application of ALD from 1 hpi onwards was highly efficacious in lowering SFV6 RNA in all tissues analysed and infectious virus levels in the blood by 24 hpi (Fig 2A,B). For mice in which infection was allowed to progress, topical ALD application resulted in a significant delay to the onset of neurological signs (Fig 2C). Consistent with this, analyses of brain tissue at post infection day 7 revealed extensive expression of SFV6-encoded mCherry throughout the brain in untreated mice, compared to treated mice (Fig 2D). Similarly, topical ALD at 1hpi post infection with the less virulent SFV4 strain, significantly reduced virus levels by 24 hpi in tissues, the blood and substantially decreased mortality, with the majority of mice surviving infection (Fig 2E-G). Importantly, protection by ALD was time-limited, as while treatment that was delayed until 5 hpi significantly lowered viral RNA levels, treatment delayed until 10 hpi did not significantly modulate the level of viral replication or dissemination by 24 hpi (Fig 2H). Together these data identify the inoculation site as a key site for viral replication during the first 24 hpi and suggests that, in untreated mice, skin IFN responses are not sufficiently robust to prevent systemic dissemination. Therapeutic targeting of this site with topically applied innate immune agonists was therefore highly effective at suppressing both the local and subsequent systemic course of infection, significantly improving survival.

**Fig. 2.**
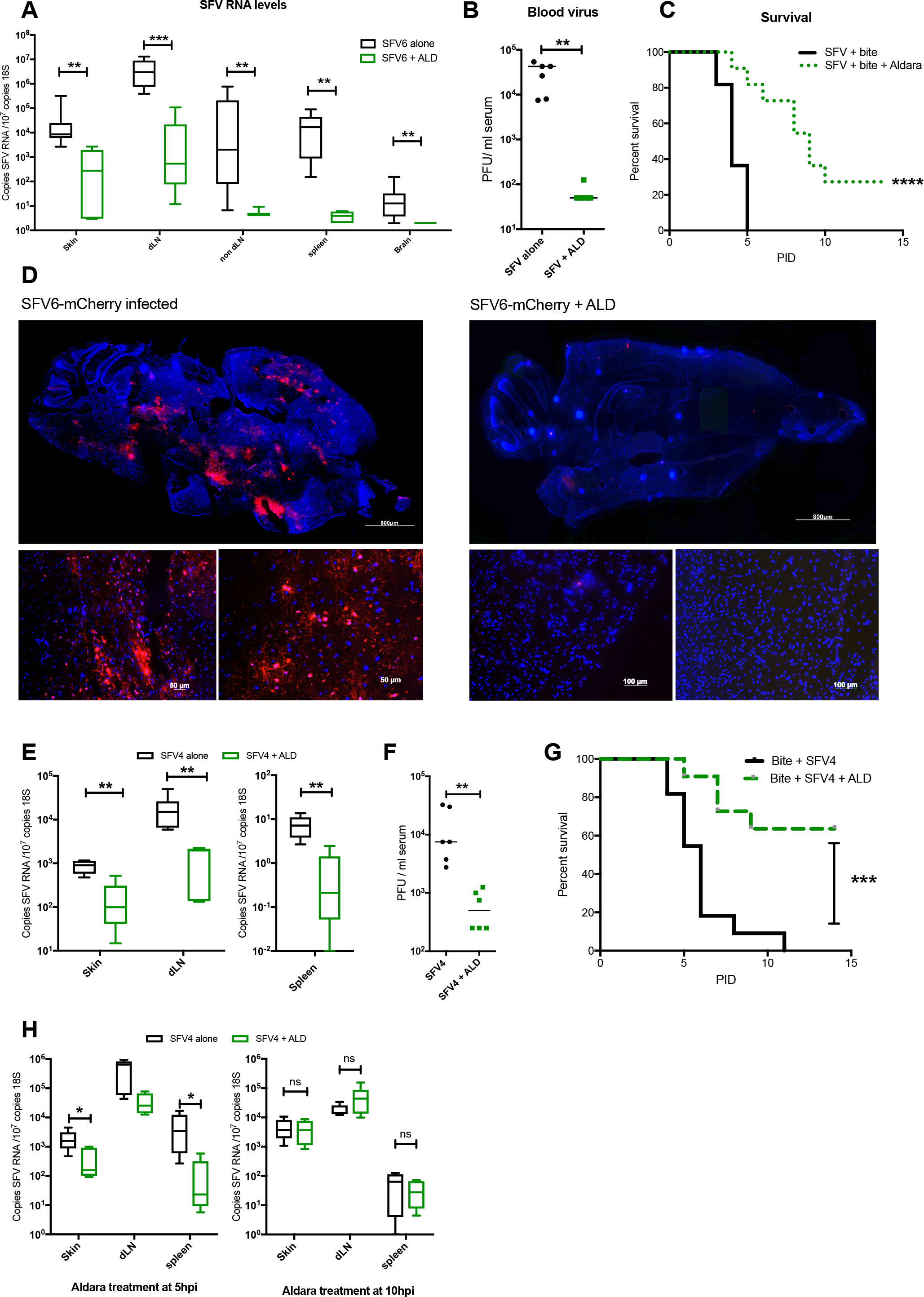
Targeted post-exposure immunomodulation suppresses the local and systemic course of infection and improves clinical outcome in mice. (A-D) Mosquito-bitten mouse skin was infected with 250 PFU SFV6 and treated with topical ALD at 1 hpi or left untreated. (A) SFV RNA and host 18S determined by qPCR at 24hpi (n=7). (B) Plaque assay of serum at 24hpi (n=7). (C) Survival of mice (n=10). (D) Mid-sagittal sections of the brain from mice infected with SFV6-mCherry stained with DAPI (blue); upper composite image was assembled from multiple photographs, while lower images show typical individual images at higher magnification. (E-G) Mice were similarly infected with SFV4 at a mosquito bite and treated with ALD or left untreated. (E) Levels of SFV RNA and host 18S were determined by qPCR at 24hpi. (F) Serum virus quantified by plaque assay (n=6). (G) Survival curve of mice (n=11) following ALD (green line) or no treatment (black line). (H) Mice were infected with SFV4 at a mosquito bite and treated with ALD at either 5 or 10 hpi. Levels of SFV RNA and host 18S were determined by qPCR at 24hpi. (n=6) *p<0.05, **p<0.01, ***p<0.001, ns=not significant (Mann-Whitney and logrank (Mantel Cox) test).

### ALD induces upregulation of cutaneous ISG via canonical type I IFN receptor signalling

Having identified the inoculation site as amenable to therapeutic targeting with topical ALD, we next wanted to determine its mode of action, as a detailed mechanistic understanding will inform the development of more targeted strategies. Importantly, ALD contains a mixture of the TLR7 agonist IMQ (5%) and isostearic acid (25%), both of which can be inflammatory in a type I IFN-independent manner (28, 29), suggesting that IFN may be dispensable for some of the beneficial effects of ALD described here. However, we found that by 24 hours post application at a mosquito bite, ALD caused widespread induction of ISG expression, with the prototypic ISGs *cxcl10, ifit1, isg15* and *rsad2* most highly expressed in the skin, most of which have been identified as key responders to alphavirus infection (17, 30). Increased expression of *cxcl10* could be detected by 16 hours and this increased significantly by 24 hours (Fig. S2E). Interestingly however, despite a clear upregulation of ISG, we could not detect any increase in type I (*ifna4, ifnb1, ifne, ifnk, ifnz*) or II (*ifng1*) IFN transcripts in whole skin biopsies, as measured by Taqman assays or custom SYBR green qPCR assays (Fig. 3A, S2A-C), suggesting ALD may activate ISG expression independent of IFN receptor signalling in the skin (*31*). Anti-viral ISG can be induced by a variety of distinct receptors, including signalling through IFN-αβ-R, IFNGR, IFNLR1 and other poorly-defined non-canonical pathways (16, 31). However, we found that activation of skin ISG was dependent on IFN-αβ-R signalling, as mice deficient in this receptor could not upregulate these prototypic ISGs in response to ALD (Fig. 3C). This was associated with a loss of protection from infection, as topical ALD had no impact on virus replication in IFN-α-R null mice (Fig. 3D,E). In addition, treatment with an ALD cream mimic that lacked IMQ but contained 25% w/w isostearic acid (28), did not protect mice from infection (Fig. 3F). Nonetheless, its likely that these excipients are required for a maximal IFN response, as injection of IMQ alone had little or no effect on ISG expression in either the skin or dLN (Figure S2D-G), nor did it have any effect on virus replication (Fig. 1E-G and Fig. S1A,B). In comparison to skin responses, ALD upregulated both IFN and ISG in the dLN; IFNγ was significantly elevated by 8h post application (Fig. S2B), while type I IFN were detectable by 24 hours, as were ISGs (Fig. 3B and Fig. S2E). In agreement with previous studies in humans (*32*), there was a limited systemic response to topical ALD, although our qPCR assay was sufficiently sensitive to detect some type I IFN expression in distal lymphoid tissue sites, but not remote skin or joints (Fig. 3G).

**Fig. 3.**
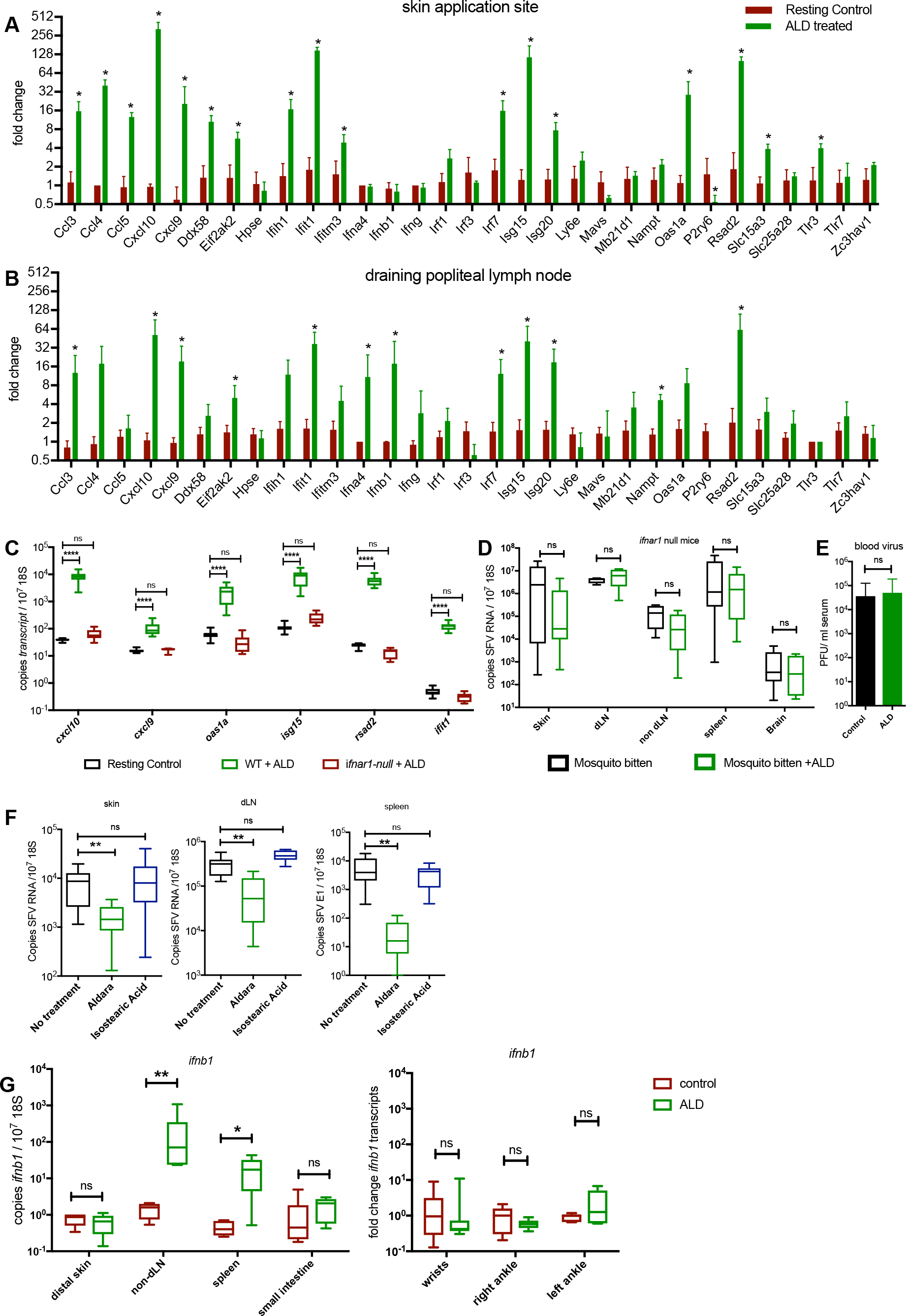
Protection by topical imiquimod is dependent on type I IFN signaling. (A,B) Mice were bitten with mosquitoes and the site treated with ALD. (A,B) Fold change gene expression of ISG in skin (A) and draining popliteal LN (B) was determined by qPCR at 24h post treatment compared to resting controls (n=4). (C) *ifnar1* −/− mice and syngeneic wild type controls were treated with a single 2mg application of topical ALD. Copy number of host ISG and 18S RNA were determined by qPCR at 24h post treatment (n=10) (D,E) *ifnar1* −/− mice (n=6) were infected with SFV6 in the presence of a mosquito bite then treated with topical ALD. (D) SFV RNA and host 18S determined by qPCR at 24hpi. (E) Serum virus quantified by plaque assay. (F) Mosquito-bitten WT mouse skin was infected with SFV6 and then treated with either topical ALD, or control cream (25% w/v isostearic acid). SFV RNA and host 18S was determined by qPCR at 24 hpi (n=6). (G) Mice were treated with topical ALD and levels of *ifnb1* determined by qPCR at 24 h (n=6) *p<0.05, **p<0.01, ****p<0.0001, ns=not significant (Mann-Whitney and Kruskal-Wallis test)).

### Skin resident cells detect topically-applied IMQ and are sufficient for mediating protection to virus

We next wanted to identify the cellular basis by which virus is detected and an anti-viral state induced and modulated by ALD at mosquito bites. Although a variety of cultured skin cells are known to express RNA-virus pattern recognition receptors, the cellular coordination of innate immune responses to arbovirus at the mosquito bite are not described (*10, 33*). We concentrated on studying cells of the dermis, as the majority of virus transmitted by mosquito is deposited here, and because epidermal cells do not express the IMQ receptor TLR7 *in vivo* (*34*). Mosquito biting results in the rapid recruitment of leukocytes including neutrophils and monocytes (9), and as shown here small numbers of pDCs (Fig 4A). To identify whether these bite-recruited leukocytes or skin-resident cells are sufficient for detecting IMQ and eliciting protection from virus, we developed a skin explant model in which freshly biopsied skin was taken from either resting skin or mosquito-bitten skin (4 h post bite, during peak leukocyte influx (9)) and infected with SFV6-Gluc *in vitro*. Infection of skin explants resulted in replication of virus, as measured by qPCR of viral RNA, functional expression of virus-encoded luciferase and release of new infectious virus into the surrounding tissue culture medium. Application of ALD to the epidermis of cultured skin explants 1 hpi resulted in a significant decrease in viral RNA, virus-encoded luciferase and a complete block in the release of new virus into the supernatant, for both explants derived from resting- and mosquito bitten-skin (Fig 4B-D). Thus, the ability of ALD to reduce viral replication occured irrespective of leukocyte recruitment triggered by infection or by mosquito bite. Furthermore, the magnitude of fold decrease in viral RNA was similar in *ex vivo* explants to that observed *in vivo* (Fig. 4E). In addition, ALD application was efficacious in limiting SFV replication when applied to non-bitten skin (Fig. 1E-H). Together this suggests that skin-resident cells were sufficient for mediating ALD-induced protection from virus and that skin-infiltrating leukocytes were not required.

**Fig. 4.**
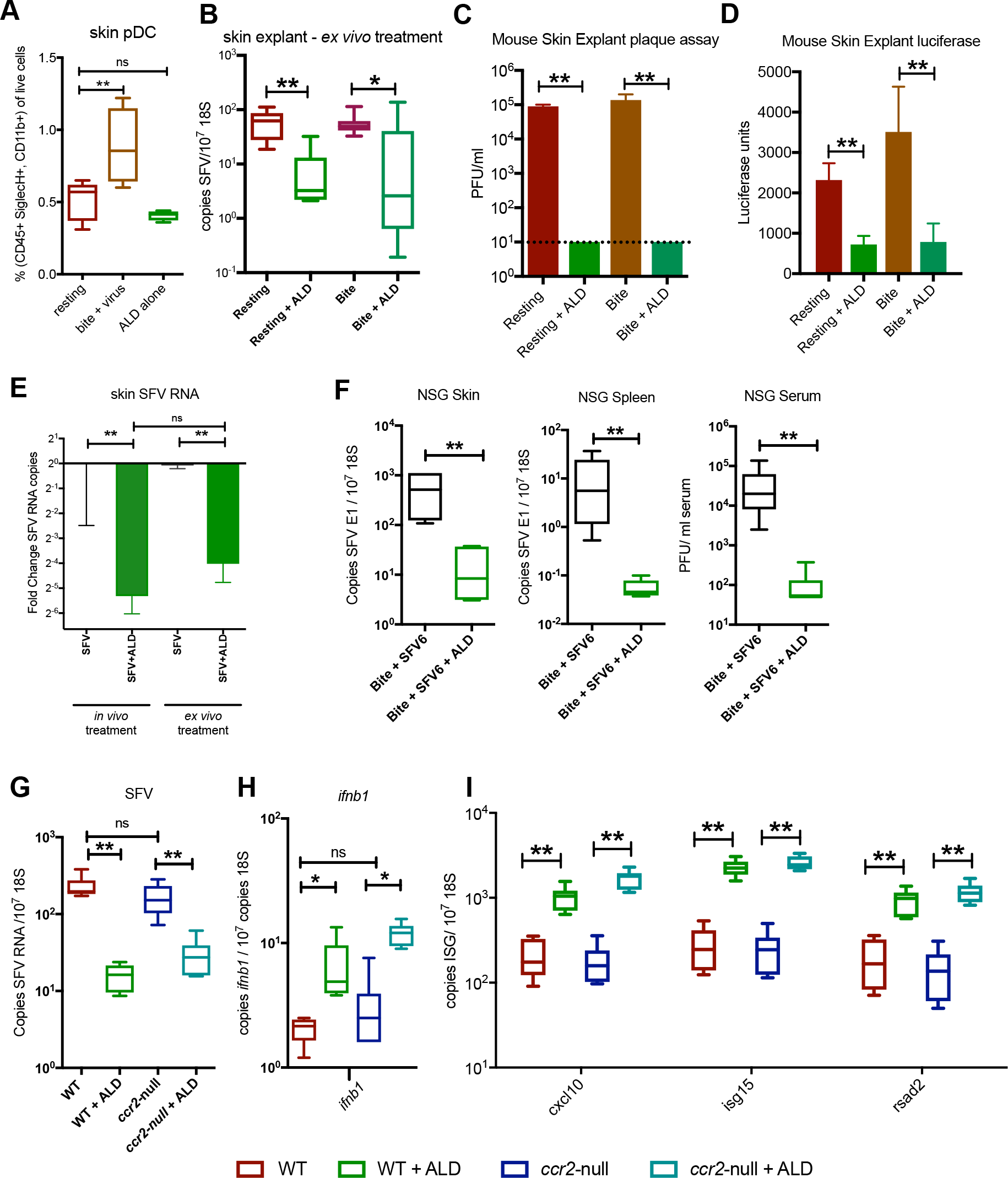
Host responses to IMQ by skin-resident cells are sufficient to mediate protection. (A) Mosquito-bitten mouse skin was infected with SFV6. At 24 hpi, skin from the inoculation site was digested to release cells and numbers of plasmacytoid dendritic cells (pDCs) quantified by FACS. (B-D) Mosquito-bitten mouse skin was biopsied at 4 hours post bite and resulting 6mm skin explants infected *ex vivo* with 10^5^ PFU SFV6-Gluc. 1 hpi explants were treated with a single application of topical ALD. (B) At 24 hpi; copy number of SFV RNA were determined by qPCR; (C) virus titers in the media quantified by plaque assay (n=6); and (D) virus-encoded Gluc assayed (n=6). (E) Mice were either infected *in vivo* at mosquito bites with or without topical ALD (at 1 hpi), or 6mm resting skin explants were infected with SFV6 *ex vivo* by needle and then treated with a single application topical ALD (at 1 hpi). Copy number of SFV RNA and host 18S was determined by qPCR at 24 hpi and compared to respective untreated infected controls (n=6) (F) Mosquito-bitten skin of NSG mice were infected *in vivo* with SFV6 and then treated with topical ALD from 1 hpi. Levels of SFV RNA was determined by qPCR in skin and spleen at 24 hpi (n=6). (G-I) 6mm skin explants from WT and *ccr2*-null mice were infected with SFV6 and treated with a single 2 mg application of topical ALD *ex vivo*. At 24 hpi, copy number of; (G) SFV RNA; (H) *ifnb1*; and (I) host ISG (*cxcl10*, *isg15*, *rsad2*) were determined by qPCR (n=6). *p<0.05, **p<0.01, ns=not significant (Mann-Whitney and Kruskal-Wallis test)

We therefore wanted to define which skin-resident cells were necessary for activating ISG expression in response to ALD. Skin-resident leukocytes include populations of γδ-T cells and ROR-γT innate lymphocytes that are necessary for some inflammatory responses to repeated ALD application (*35*). However, we found that NOD SCID Gamma (NSG) mice, which lack all functional lymphocytes and innate lymphoid cells, were similarly protected from infection at 24hpi by ALD (Fig. 4F). This suggests early skin responses to ALD are activated by either stromal cells or non-lymphoid leukocytes, of which myelomonocytic cells are the most populous in the dermis. Skin dermis-resident myelomonocytic cells are derived from either bone marrow (BM) precursors (requiring expression of the chemokine receptor CCR2 and includes dermal DCs, monocytic cells and some macrophage sub-populations); or are derived from non-BM embryonic sites of haematopoiesis (not requiring CCR2 expression and limited to a sub-population of dermal macrophages) (*36*). To determine if BM-derived skin resident myelomonocytic cells are required to mediate ALD-protection, we infected skin explants from *ccr2*-null and WT mice and treated them with topical ALD. ALD was similarly efficacious irrespective of *ccr2* status, suggesting that the vast majority of skin-resident dermal DC and monocytic-derived cells are not required (Fig. 4G). Indeed, IFN and ISG induction in response to ALD application was similar in *ccr2*-null skin as compared to WT skin (Fig. 4H,I), suggesting that ALD-responding cells were skin-resident cells not derived from myeloid BM-precursors.

### Skin macrophages and dendritic cells communicate with stromal cells to mediate ALD-induced protection from virus

Together, the above data suggests that the skin-resident cell type responding to ALD was either a population of non-BM derived macrophages, mast cells, stromal cells or a combination of these. It is not possible to deplete all these cell types; therefore, we additionally undertook a comprehensive approach to define responses of all these cell populations. This involved analysing the gene expression profile of skin inoculation-site cells isolated by FACS, to reveal the cell-specific basis for innate immune response to virus and separately to ALD (Fig. 5).

**Fig. 5.**
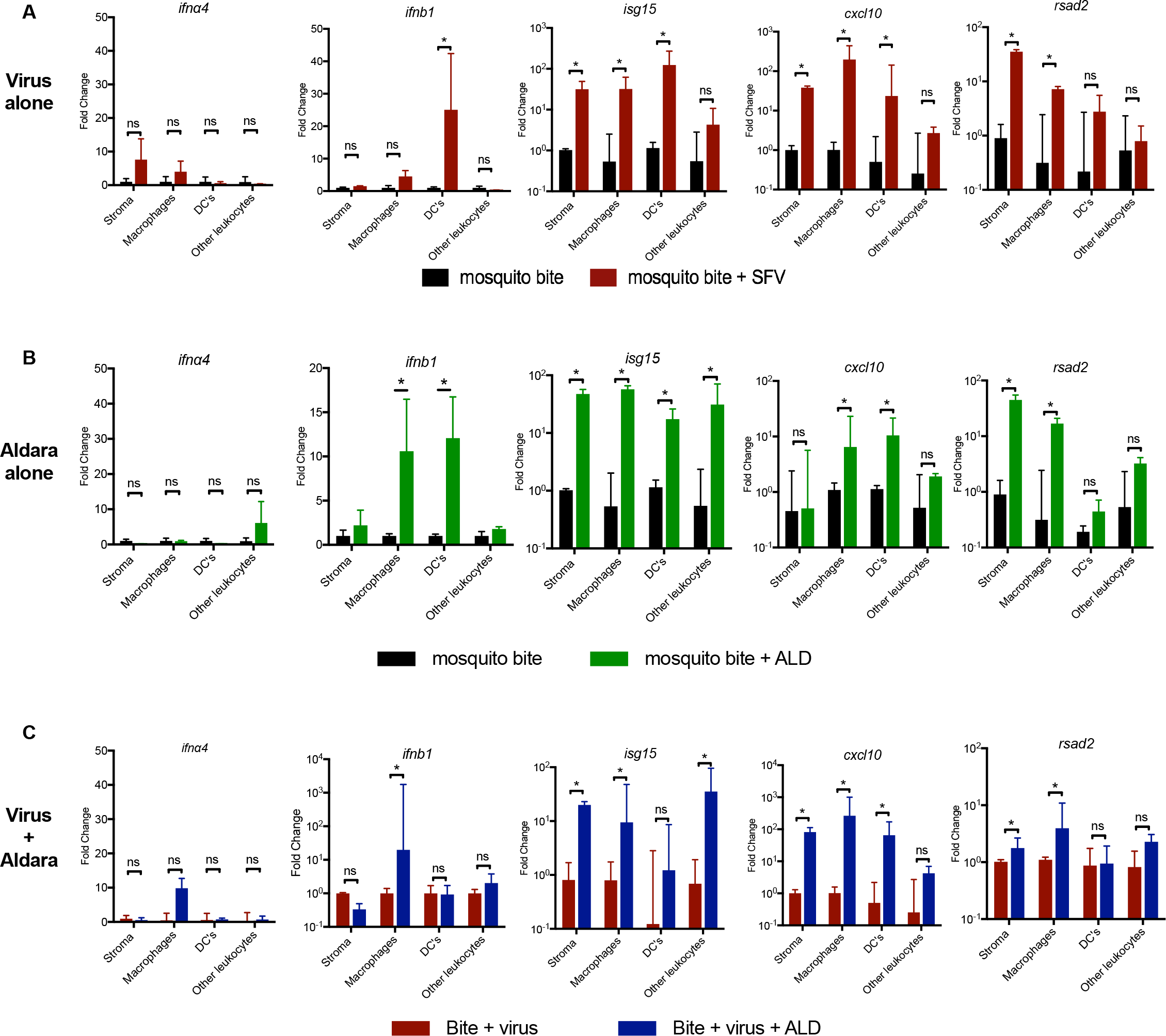
Topical IMQ targets dermal macrophages and dendritic cells to activate tissue-wide ISG expression. Mosquito-bitten mouse skin was digested to release cells and then FACS-sorted at 4°C into “stroma”, “macrophage”, dendritic cell “DC’s” and “other leukocyte” compartments as a bulk sort. All steps were undertaken in presence of transcriptional inhibitors. Copy number of gene transcripts were determined by qPCR for; *ifna4*, *ifnb1*, *isg15*, *cxcl10*, *rsad2* and 18S RNA was (n=4). Cells were sourced from mosquito bitten skin that was either; (A) infected with SFV6 alone; (B) treated with topical ALD alone; (C) infected with SFV6 and treated with ALD. *p<0.05, ns=not significant (Mann-Whitney)

Although skin-resident cells are sufficient to mediate IMQ-responses, this does not necessarily exclude a role for mosquito-bite elicited responses at later stages. For this reason, and because arboviruses are all naturally transmitted into skin bitten by arthropods, we studied cellular responses to virus and ALD in mosquito-bitten skin. We adapted a previously-defined FACS gating strategy (*36*), to isolate populations of; macrophages; dendritic cells; stromal cells; and all other CD45+ leukocytes as a bulk population (Fig. S3A,B). These isolated populations were then interrogated for their expression of *ifna4, ifnb1, isg15* and *rsad2* because they have been previously implicated as key for host responses to alphaviruses (*30, 37, 38*) and also the prototypic ISG *cxcl10*.

The cellular coordination of innate immune responses to virus at the tissue-wide level in mosquito-bitten skin are poorly defined. Therefore, we firstly analysed responses of skin cells to virus infection alone in the absence of ALD (Fig 5A). At 24hpi, sorted stromal cells and macrophages were the only cells that expressed high levels of virus structural gene transcript E1, indicative of active replication, while DCs expressed only very low levels of viral RNA (Fig. S3C). This, along with our previous studies (9), suggest that stromal cells and macrophages are key targets for viral replication of SFV at mosquito bites *in vivo*. Importantly, the only cell type expressing type I IFN in response to virus were DCs, which was limited to *ifnb1* only, as no *ifna4* was detected. Thus, DCs were the primary initiators of anti-viral ISG responses at virus-infected mosquito bites. However, dermal DC were present at low numbers in the skin at 24hpi, suggesting their total contribution to the tissue-wide IFN response was minimal (Fig. S3B). This agrees with our whole tissue analysis that demonstrated little IFN expression before 24 hpi (Fig. 1A). Thus, although stromal cells and macrophages exhibited high levels of viral RNA they did not make detectable type I IFN. Nonetheless they expressed some ISG including *rsad2, cxcl10 and isg15* by 24hpi (Fig. 5A), presumably in response to DC-expressed IFN cues. All other leukocytes included in the bulk sort did not express IFN or upregulate ISG in response to virus.

Next, to determine which cells were activated by ALD, mosquito-bitten mouse skin was treated with ALD alone (Fig. 5B, Fig. S3E). Crucially, in contrast to virus infection, ALD induced *ifnb1* expression in both the more populous skin-resident macrophages and dermal DC, while stromal cells and all other leukocytes lacked type I IFN transcripts. There was also a trend for higher *ifna4* in macrophages, although this was too variable to be statistically significant in our experiments. In response, stromal cells upregulated *rsad2* and *isg15* to high levels. As viruses have evolved mechanisms to antagonise IFN signaling we next wanted to determine if virus infection modulated this response to topical ALD. Therefore, we infected mosquito bites with virus, either with or without ALD application at 1 hpi and assayed type I IFN and ISG expression at 24 hpi in each cell type (Fig 5C). Despite the lower titres of virus in ALD-treated skin by 24 hpi (and therefore less activation of IFN signalling via virus sensing), we found equal or higher fold increases in the expression of both type I IFN and ISG in the macrophage population. ISG upregulation was also more robust in macrophages, DC and stromal cells, with the exception of *rsad2* in DC, suggesting that ALD-induced upregulation during virus infection was ISG and cell type-specific. Activation of higher IFN expression in macrophages was important for two reasons; firstly, macrophages were significantly more numerous in the skin than dermal DC at 24 hpi (Fig. S3B) and secondly, they constituted an important source of virus replication in the first 24 hours of infection (Fig S3C and (9)). ALD-elicited IFN appeard to be functional, as type I IFN negative stromal cells exhibited a significant decrease in viral RNA with ALD treatment by 24hpi, as did virus-infected macrophages (Fig. S3C).

Together, this shows that ALD upregulated type I IFN in skin-resident macrophages that, in addition to responses of the less frequent dermal DCs, acted to induce ISG expression and thereby reduce viral replication in the skin. Furthermore, because dermal DC-deficient *ccr2*-null skin (*36*) was not compromised in its ability to express ISG or reduce virus titres in response to ALD (Fig. 4H,I), we suggest that skin-resident macrophages alone are sufficient to mediate protection. Thus, responses by skin-resident macrophages to ALD are likely sufficient to confer protection against virus infection.

#### Stromal cells integrate cues from IMQ-treated leukocytes to resist infection with virus

We next wanted to determine whether signals from IMQ-activated macrophages and DCs are alone sufficient to confer protection on stromal cells from virus infection and whether this required cell-to-cell contact. The majority of dermal stromal cells are either fibroblasts or keratinocytes, both of which can be experimentally-infected with a range of arboviruses (*10, 39, 40*), including SFV4 (Fig. 6A-D). Some cultured fibroblasts have been described to express a variety of receptors that sense virus, including the IMQ receptor TLR7 (*41*). However, we found that FACS-isolated stromal cells *ex vivo* lacked detectable TLR7 (Fig. S3D). In addition, our cultured primary dermal fibroblasts (Fig. 6B) and keratinocytes (Fig. 6E) did not show any response to IMQ following SFV infection; IMQ did not protect cells from infection; ISG *cxcl10* expression was not induced (even at high doses, Fig. 6C); and media from fibroblasts pre-treated with IMQ for 24 hours did not confer protection to other stromal cells from subsequent virus infection (Fig. 6D). Thus, both our *in vivo* (Fig. 5B) and *in vitro* analyses suggest stromal cells were not able to respond to IMQ alone.

**Fig. 6.**
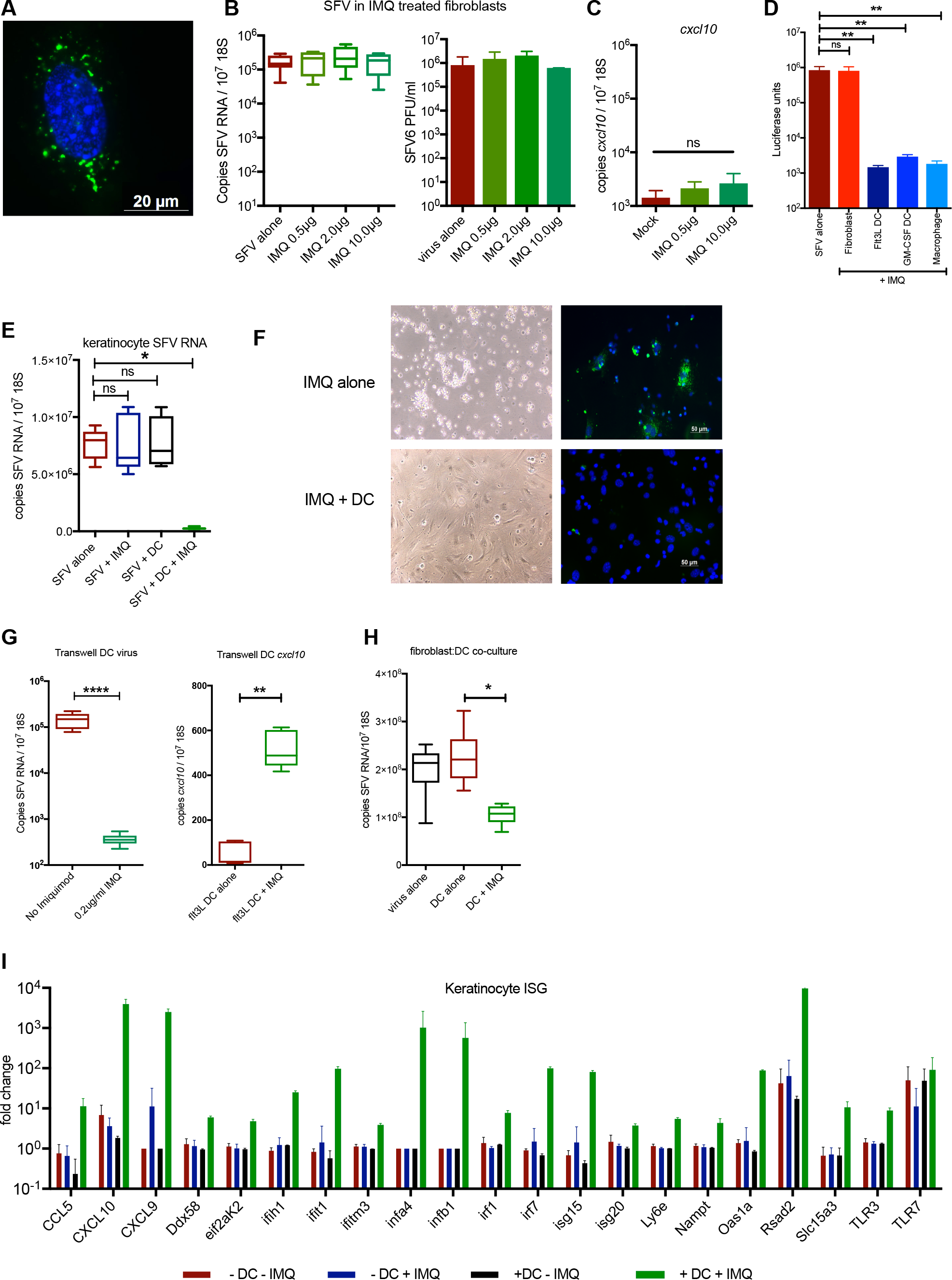
Protection of skin stromal cells from virus by IMQ requires signals from myeloid cells. (A) Primary keratinocytes were infected with SFV4(Xho)-EGFP. At 6 hpi replication complexes (green) were present throughout cytoplasm (DAPI, blue). (B,C) Primary murine embryonic fibroblasts were infected with SFV6 in vitro at an MOI of 0.1. At 1hpi, cells were treated with either 0.5, 2 or 10 μg/ml IMQ and at 24 hpi; (B) copy number of SFV RNA determined by qPCR (n=6) and infectious virus in the media were quantified by plaque assay; and (C) copy number of host *cxcl10* gene transcripts was determined by qPCR (n=6). (D) Primary dermal fibroblasts and BM-derived Flt3L DCs, GM-CSF DCs and M-CSF macrophages, were treated with 0.5 μg/ml IMQ for 24 hours. The resulting conditioned media was aspirated and placed on SFV6-Gluc-infected fibroblasts (1 hpi with at an MOI of 0.01). Virus-encoded Gluc was assayed at 24 hpi (n=6). (E-G) Primary keratinocytes were infected *in vitro* with SFV6 (E,G) or SFV4(Xho)-eGFP (F) at an MOI of 0.1, then at 1 hpi treated with 0.5 μg/ml IMQ in the presence or absence of Flt3-L DC’s separated by a 0.5 μm transwell membrane. (E) Copy number of SFV RNA and host 18S RNA in the keratinocytes were determined by qPCR at 24 hpi (n=6). (F) SFV-encoded eGFP shown as green, with cell nuclei counter-stained with DAPI (blue). (G) Copy number of SFV RNA and *cxcl10* transcripts were determined in DC by qPCR at 24hpi (n=6). (H,I) Primary fibroblasts (H) or keratinocytes (I) were infected *in vitro* with SFV6 at an MOI of 0.1 in the presence or absence of Flt3L-derived DCs separated from stromal cells by a transwell membrane. At 1 hpi cells were either left untreated or given 0.5 μg/ml IMQ. (H) Copy number of SFV RNA were determined by qPCR at 24hpi (n=6). (I) Gene expression of keratinocyte ISG were determined by qPCR (n=4). *p<0.05, **p<0.01, ****p<0.0001, ns=not significant (Mann-Whitney and Kruskal-Wallis test)

To determine if soluble factors from IMQ-stimulated DCs or macrophages could influence the susceptibility of primary cultures of skin fibroblasts to infection, primary cultures of leukocytes were stimulated with IMQ for 24 hours and their supernatant given to fibroblasts at 1 hpi with luciferase-expressing SFV6-Gluc (Fig. 6D). While supernatant from resting leukocytes had no effect on fibroblast susceptibility to infection, tissue culture supernatant from DC or macrophages treated with IMQ protected fibroblasts from infection, as measured by virus-encoded luciferase.

As infected myeloid cells and DC are susceptible to arbovirus-encoded anti-IFN mechanisms (*42, 43*), we next determined if infected DC were still able to provide IMQ-induced protection and trigger an anti-viral state in stromal cells. To do this we used a transwell system in which DC were separated from stromal cells by a cell-impermeable membrane. All cells were infected with SFV4(Xho)-EGFP and treated at 1 hpi with either 0.2 μg/ml IMQ or saline control (Fig. 6E-H). DC cultures themselves became infected with SFV, exhibiting high level expression of viral RNA (Fig 6G). Nonetheless, DC were able to resist infection upon addition (1hpi) of IMQ and consequently expressed higher levels of the ISG *cxcl10*. Importantly, while the addition of IMQ alone, or DC alone, had no effect on the ability of co-cultured stromal cells to resist infection, the combined presence of both IMQ and DC did protect keratinocytes (Fig. 6E,F and Fig S4) and fibroblasts (Fig. 6H). This was evident by a significant decrease in viral RNA expression (Fig 6E,H) and by microscopic inspection of the stromal cell monolayer that otherwise became decimated by 24hpi, with the remaining intact cells also positive for virus-encoded eGFP (Fig. 6F, Fig. S4). Protection from virus in stromal cells occurred in conjunction with the induction of stromal ISG expression (Fig. 6I). While keratinocytes did not express ISG or type I IFN in response to either virus alone, IMQ alone, or both; they did significantly increase ISG expression with the addition of DC to the transwell insert. Thus, IMQ-induced stromal cell protection from virus occurs in conjunction with the induction of ISG expression that was licenced by extracellular cues derived from IMQ-responsive leukocytes.

#### ALD protects mice and human skin from infection with a variety of genetically-distinct medically-important viral pathogens

We finally wanted to determine whether activating IFN pathways by ALD at the skin inoculation site could have potential applicability to a variety of genetically-distinct arboviruses of medical importance. Arboviruses are derived from several evolutionary- and genetically-divergent groups including; Togaviridae, Flaviviridae and Bunyavirales. We therefore analysed the ability of ALD to modulate the outcome of infection to a relevant representative of each of these distinct groups in mice and human skin explants. The prototypic Bunyamwera virus (BUNV, genus Orthobunyavirus, family Peribunyaviridae, order Bunyavirales), which like SFV is also transmitted by *Aedes* mosquitoes, has well-described potent IFN antagonism (4) and therefore may be resistant to ALD treatment. However, when mosquito-bitten BUNV-infected inoculation site was treated with topical ALD at 1hpi, there was a significant decrease in viral RNA in target tissues and blood (Fig. 7A-C). Similarly, we determined if topical ALD could prevent dissemination of CHIKV (a medically important pathogen of the Togaviridae family that has caused widespread outbreaks of disease) to mouse joints that were remote from the inoculation site (Fig. 7D-G). Mice treated with ALD 1 hpi exhibited significantly lower levels of CHIKV RNA (qPCR) and infectious virus in the ankle joint contralateral to infection, and also a decrease in CHIKV RNA in the wrists of the forelimbs. Finally, to determine if ALD could also reduce viral replication in human skin, we cultured explants of freshly biopsied human skin, infected them with virus and treated with topical ALD to the epidermis at 1 hpi (Fig. 7H,I). Application of ALD resulted in a significant decrease in viral replication for both CHIKV and ZIKV (genus Flavivirus of the family Flaviviridae), as measured by viral RNA and also infectious virus for ZIKV. Together these studies demonstrate that an innate immune agonist that primarily targets inoculation-site skin-resident macrophages, has a significant post-exposure prophylactic effect on the replication and dissemination of a variety of genetically-distinct medically-important viruses.

**Fig. 7.**
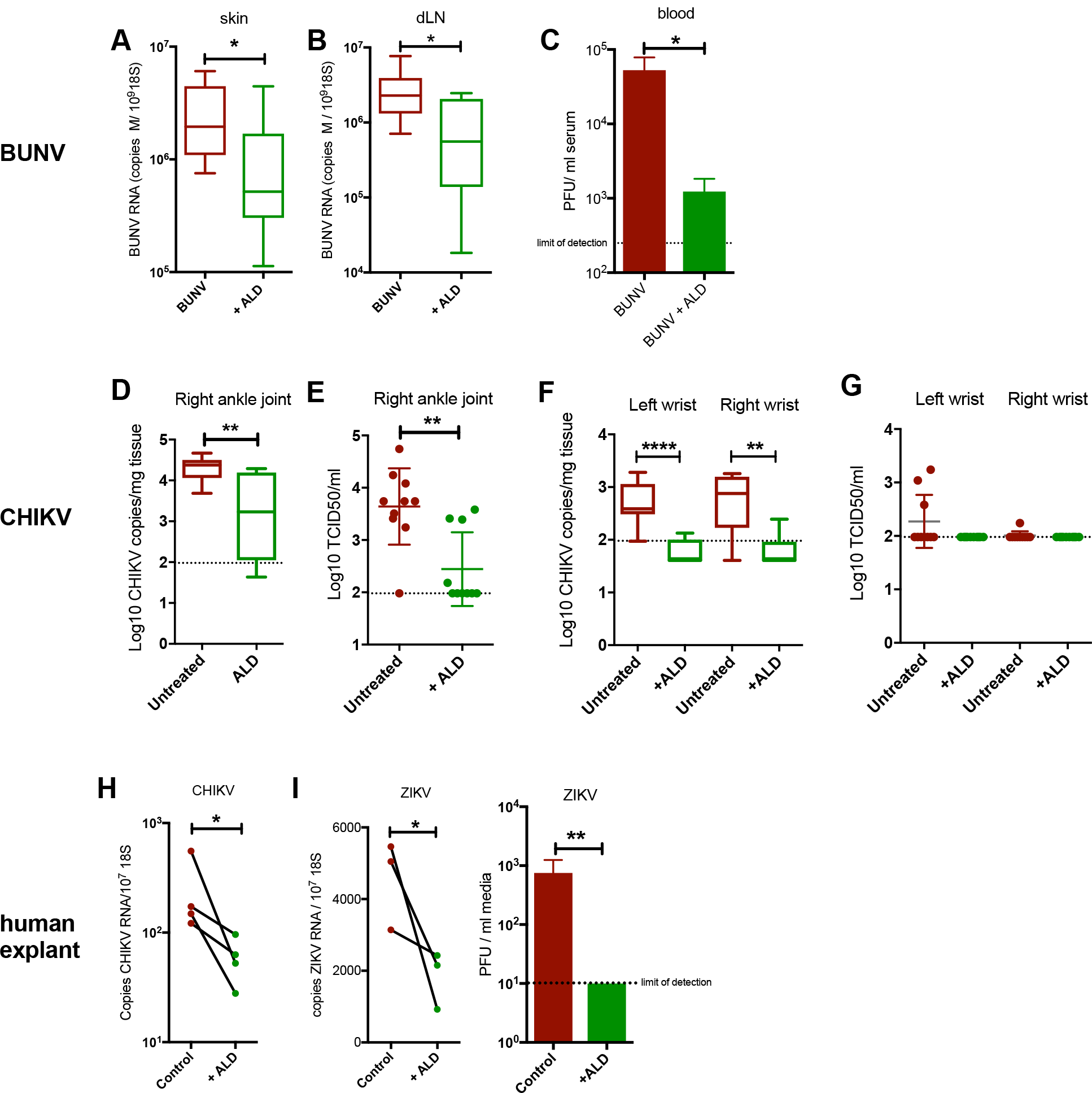
Topical IMQ protects both mouse and human skin from varied, genetically-distinct arboviral threats. (A-C) Mosquito bitten skin of mice (n=6) were infected with BUNV and treated with topical ALD from 1 hpi. At 24 hpi, copy number of BUNV RNA was determined by qPCR in skin (A) and dLN (B); and infectious virus in blood determined by plaque assay (C). (D-G) Mice were infected with 10^6^ PFU of CHIKV (s.c.) and treated with topical ALD from 1hpi onwards. Dissemination of virus to distal joints were assessed at 24hpi in the right ankle joint. (D,E) and both wrist joints (F,G). Levels of CHIKV RNA were determined by qPCR and infectious virus titers quantified by TCID50 (n=10). (H,I) Bisected human skin biopsies were cultured as explants and infected with virus (1×10^5^ PFU CHIKV (H) or 0.5×10^5^ ZIKV (I)). One half of each biopsy was treated with a single application (2 mg) of topical ALD *ex vivo* 1hpi; while the other half was left untreated. Levels of CHIKV and ZIKV RNA was determined by qPCR at 24 hpi. For ZIKV infection, virus titers in the media were also quantified by plaque assay (n=3). *p<0.05, **p<0.01, ****p<0.0001, ns=not significant (Mann-Whitney and paired Student’s t-test for H and I).

## Discussion

We demonstrate that innate immune responses to virus in the skin is a limiting factor for suppressing subsequent systemic infection. Critically, we show that therapeutic modulation of these pathways in the first hours of infection can suppress disease and improve clinical outcome in mice. This work therefore identifies the inoculation site as an important site of arboviral replication and that skin innate immune virus-sensing pathways must be sufficiently robust to act as determinants of the clinical course. Therapies that either target IFN induction, or alleviate the virus-encoded mechanisms that inhibit IFN, may therefore be efficacious for human infection. Targeting a common pathway at the inoculation site, as shown here to suppress infection by several genetically-distinct arboviruses, may circumvent the need to develop multiple species-specific antivirals, as their development and future use remains challenging due to the unpredictable nature of arbovirus outbreaks and difficulty associated with reaching an accurate and timely clinical diagnosis.

In this study, we identified the key skin cell types in mice that coordinate anti-viral immune responses to virus and those that respond to topical ALD. We found that skin type I IFN expression to virus was restricted to the relatively infrequent DC population. In contrast, macrophages and stromal cells could not elicit IFN expression, despite a high level of viral RNA in these cells. Thus, skin-resident DC are key activators of the type I IFN response at the inoculation site to virus. Such specificity may reflect either; cell-type specific susceptibility to virus-encoded mechanisms to antagonise IFN activation; or the particular cell-tropism of our model arbovirus, SFV.

We suggest that the observed efficacy of our exemplar immune-modulator, ALD, was explained by activation of dermal macrophages. These cells were not only otherwise deficient in their IFN expression in response to virus alone, but also can replicate high levels of virus (9). Therapies that specifically target dermal macrophages may provide yet enhanced efficacy.

ALD was no longer efficacious where treatment was delayed for 10 hours in our mouse model. We suggest that this timing most likely reflects specific aspects of our model system that supports rapid dissemination of SFV, itself a mouse-adapted strain. It is not clear how these kinetics apply to human infection by mosquito, although it is widely-accepted that virus may take considerably longer (several days) to disseminate from human skin to the blood (*44*). This in turn may mean that the window available for post-exposure intervention in humans is conveniently longer than that observed in our model, although virus-, host- and environment-specific factors are likely to define its length. However, because *Aedes* sp. mosquitoes are day-biting and their bites are typically visible within minutes, it is possible that there is a sufficient time-window for post-exposure prophylactic treatment between infection/biting and awareness of a bite/application of cream in humans.

As such, we suggest that therapeutic intervention at the inoculation site represents a novel strategy for targeting arbovirus infection that merits further investigation. We suggest that with refinement, a modified form of our exemplar immune-modulator (topical ALD) may have potential as a novel treatment strategy for this globally-important disease category. Thus, further work that refines the most efficacious method for targeting IFN induction in macrophages, opens up the possibility of broadly applicable, likely cost-effective, therapeutic interventions for arbovirus infection.

We maintained application of ALD for only five hours to a single site, suggesting systemic side effects are less likely than with prolonged exposure. However, effective post-exposure prophylaxis might entail treatment of multiple, discrete, suspected arbovirus-infected mosquito bites, even with e.g. vector repellent strategies employed. Thus, it will be important to refine any future formulations such as to render the application of treatment practical. However, it should be noted that e.g. ALD has been widely used for many years for a variety of dermatological conditions, including in primary care, and is generally well-tolerated by patients, even when repeatedly applied at the same site for several weeks (*45*). Additionally, our approach would likely entail topical immunomodulation over a smaller surface area than those typically treated with ALD for its licenced indications. Finally, it is noteworthy that outbreaks of arbovirus infection are explosive in nature and can be highly seasonal, enabling possible use of any prophylactic strategies to be more intensively promoted when risk of infection is known to be high. Furthermore, it is possible that those patients deemed at higher risk of complications following arbovirus infection, such as immunosuppressed individuals, or e.g. those that are more susceptible to severe dengue, may particularly benefit from this strategy.

## Materials and Methods

### Study design

This controlled laboratory study was initiated to determine whether anti-viral innate immune response to arbovirus at the mosquito bite can be therapeutically-manipulated to suppress the subsequent systemic course and improve outcome of infection in mice. We investigated mammalian host responses to representative viruses from three genetically-divergent groups of arboviruses. Two of the viruses (ZIKV and CHIKV) are medically important emerging arboviruses, while the others (SFV and BUNV) are model viruses that replicate efficiently in immune competent mice. As previously described (9), *Ae. aegypti* bite sites were infected with mosquito cell culture-derived virus in a sub-μl volume by hyperfine needle. The inclusion of a mosquito bite in our animal model was crucial, as the resulting inflammatory milieu in the skin; increases edema; alters skin leukocyte composition; modulates dissemination of virus from skin; and as a consequence, host susceptibility to infection. *Ae. aegypti* mosquitoes were used as they are responsible for transmitting the greatest number of arbovirus infections worldwide and are the principle vector for ZIKV, DENV and CHIKV. Infected mosquitoes were not used to inoculate virus to mice as the inoculum supplied by biting mosquitoes was too variable and unpredictable to allow effective comparisons.

Innate immune responses to virus at the infected mosquito bite are not well understood. Our initial observation that susceptibility to SFV infection could be significantly decreased by topical application with ALD, led us to focus subsequent experiments on determining its mechanism of action and relevance to infection with other viruses. On the basis that immune responses in mice may not always resemble those in humans, we also investigated the ability of ALD to suppress infection with the human pathogens ZIKV and CHIKV in human skin explants. Sample sizes and end points were selected on the basis of our published experience with arbovirus infection. Wherever possible, preliminary mouse experiments were performed to determine requirements for sample size, considering the available resources and ethical use of animals. Animals (gender and age-matched) were assigned randomly to experimental groups. For plaque assays, luciferase assays and qPCR, samples were coded prior to analysis, to limit bias. For qPCR, each result represents the median of 3 or 4 technical replicates of one biological replicate. For plaque assay, viral stocks and biological samples was serially diluted and each dilution assayed in duplicate. Biological replicates from mice were excluded from analysis if injection of virus inadvertently punctured a blood vessel. No outliers were removed from these studies.

## Administration of innate immune agonists

Anesthetized mice were injected with either 6μg poly(I:C), poly(dA:dT) and IMQ (Invivogen) into skin as a 4μl aqueous volume using Hamilton syringes, at the same site as virus infection (using upper skin of left foot (*9*)). Poly (I:C) and poly (dA:dT) were purchased pre-complexed with transfection reagent Lyovec (as the receptors for these agonists are cytoplasmic and widely expressed). 2mg Aldara (5% w/w IMQ; 25% w/w isostearic acid) was applied topically to the site of virus inoculation. Because Aldara (ALD) is a cream that can become removed by cage bedding, it was re-applied once at 6 hpi to maintain dosing. Although it is difficult to define the exact dose of IMQ provided using topical application of cream, previous studies have shown that dosing results in limited systemic absorption, with IMQ concentration peaking at 0.4 ng/ml blood in humans (*32*), with most being retained in the absorbent stratum corneum (the outer dead keratinized layer) or otherwise removed by external processes.

## Cell culture, viruses and mice

BHK-21 cells and C6/36 mosquito cells were grown and SFV4, SFV6 and BUNV prepared, as previously described (9). Mouse leukocytes were derived from BM-precursors; using 10ng/ml M-CSF for 6 days (macrophages); 20ng/ml GM-CSF for 6 days or 200 ng/ml Flt3L for 10 days (dendritic cells). C57Bl/6-deived primary keratinocytes (Cell biologics, USA) were grown in complete epithelia cell media as per manufacturer’s instructions. Skin fibroblasts were derived from adult C57Bl/6 dermis, digested with collagenases, dispase and DNAse to release cells and grown for 12 days in complete DMEM.

The pCMV-SFV4 and pCMV-SFV6 backbone for production of SFV has been previously described (*26, 46*). The EGFP marker gene was inserted into the C-terminal region of nsP3 via a naturally-occurring XhoI site. mCherry and Gaussia luciferase (Gluc) were separately inserted as a cassette under the control of a duplicated sub-genomic protomer 3’ with resulting viruses referred to here as SFV6-mCherry and SFV6-Gluc (*47*). Plasmids were electroporated into BHK cells to generate infectious virus. ZIKV was derived from a Recife isolate (ZIKV PE243), kindly supplied by Prof Alain Kohl (MRC-University of Glasgow Centre for Virus research). In CHIKV mouse experiments, CHIKV Indian Ocean strain 899 (<u>FJ959103.1</u>) was used (Prof. C. Drosten, University of Bonn, Germany) (*48*). For human explant studies, CHIKV was made from the infectious clone derived from the LR2006_OPY1 isolate (DQ443544). Viruses were grown once in BHK-21 cells, then passaged once in C6/36 cells and titrated prior to use. Unless otherwise specified, all mice were 6-8-week-old wild type mice (C57bl/6J). All mice were derived from a locally bred-colony maintained in a pathogen-free facility, in filter-topped cages and maintained in accordance with local and governmental regulations. Ccr2-deficient mice were originally obtained from the Jackson Laboratory (stock number 004999).

## Mosquito biting and virus infection

To ensure mosquitoes bit a defined area of skin (upper side of the left foot), anesthetized mice were placed onto a mosquito cage containing *Ae. aegypti* mosquitoes (9). Bitten skin was injected with virus in a 1 μl volume into the skin using either; 250 PFU SFV6, 1×10^4 PFU SFV4, or 2.5×10^4 PFU BUNV. For survival curves, mice were monitored closely and culled when they reached clinically defined end-points of disease. For CHIKV infection of mice, 3-4 week-old mice were infected by s.c. injection of 10^6^ PFU of CHIKV in the hind-left foot pad. For skin explant studies, tissue was dissected to remove sub-dermal tissues, to leave dermis exposed, and immediately infected with by placing lower dermis into virus-containing solution for one hour. Explants were then washed in saline and cultured in complete DMEM media at 37°C/5% CO_2_. ALD was applied topically to the epidermis at 1 hpi. For human explants, informed consent was obtained from the volunteer donors, and skin sourced from areas that were relatively protected from environmental insult and had no obvious lesions (upper inner arm). Human studies were performed following ethical approval and in accordance to all applicable regulations. All further details of reporter viruses, cells, primers and mice used can be obtained from the authors.

## Gene expression analysis and flow cytometry

Viral RNA and hosts gene transcripts were quantified by qRT-PCR and infectious virus by end-point titration, as described previously (*48*). qPCR primers for SFV and CHIKV amplified a section of E1 and primers for BUNV targeted segment M (9), while primers for ZIKV amplified a section of the env gene. For SFV, CHIKV and ZIKV, qPCR assayed measured the sum value of both genome and sub-genomic RNA. For FACS, skin tissue samples were enzymatically digested with collagenase, dispase and DNAse, stained with antibodies and a viability dye (9). Cells were analysed on a CytoFLEX (Beckman Coulter Life Sciences). For cell sorting, samples were kept on ice with transcriptional inhibitors and sorted using an Influx cell sorter (BD).

## Immunohistochemistry and histology

Tissues were fixed in 4% methanol-free paraformaldehyde (Thermo Scientific) then dehydrated in an increasing concentration of sucrose. Tissue was embedded in Optimal Cutting Temperature (OCT) compound (Agar Scientific) and sectioned. Tissue sections were stained with DAPI mounting media and imaged on a Zeiss Axioskop.

## Statistical analysis

Data were analyzed using Prism Version 7 software. Levels of viral RNA and infectious titres from virus-infected mice were not normally distributed and were accordingly analysed using the non-parametric based tests Mann-Whitney or Kruskal-Wallis test with Dunn’s multiple comparison test where appropriate, unless otherwise stated in figure legends. All such column plots show the median value +/− interquartile range. Where data were normally distributed, data were analysed using ANOVA with Holm-Sidak’s multiple comparison test and plotted with mean values. Survival curves were analyzed using the logrank (Mantel Cox) test. All plots have statistical significance indicated; *p<0.05, **p<0.01, ***p<0.001, ****p<0.0001, ns=not significant.

## Supplementary Materials

Fig. S1. Pre-exposure of skin to topical ALD increased host resistance to infection with SFV4.

Fig. S2. Type I IFN and ISG expression in skin and draining LN following IMQ application.

Fig. S3. Gene expression analysis of skin inoculation site-derived FACS isolated cells

Figure S4. IMQ-mediated protection against virus infection in keratinocytes is dependent on help from leukocytes.

## Acknowledgments

We thank Carolien De Keyzer for assistance with the CHIKV mice studies; the University of Leeds Faculty of Medicine & Health Flow Cytometry and Imaging Facilities; the University of Leeds St James’ Biomedical Services for assistance with mouse studies.

