## Supplemental material for "Pan-viral protection against arboviruses by targeting inoculation site-based skin macrophages"

**Fig. S1**

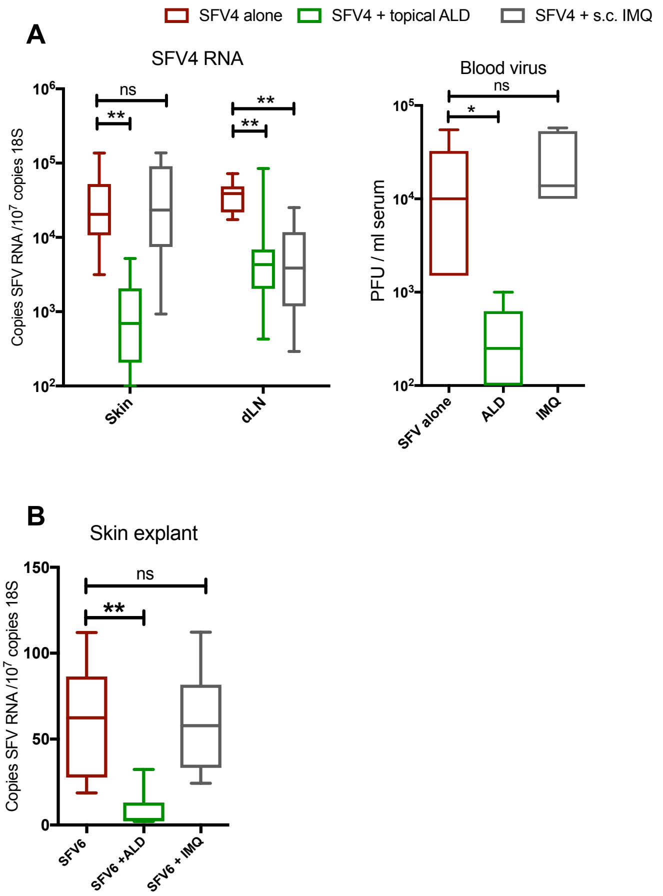


**Fig. S1. Pre-exposure of skin to topical ALD increased host resistance to infection with SFV4.**

Mice were treated with either 2 mg ALD (5% IMQ) applied topically, or to 6 µg of IMQ injected subcutaneously s.c. 1h prior to infection with 10,000 PFU SFV4 at the same site.
(A) Copy number of SFV RNA (E1 gene) and host 18S was determined by qPCR at 24 hpi (n=6). Virus titers in the serum were quantified by plaque assay (n=6).
(B) 6mm skin explants from mice were infected with 100,000 PFU SFV6 and treated with a single application of topical ALD or 6µg of IMQ *ex vivo*. Levels of SFV RNA and host 18S were determined by qPCR at 24hpi. (n=6)
*p<0.05, **p<0.01, ***p<0.001, ns=not significant (Kruskal-Wallis test)

**Fig. S2**

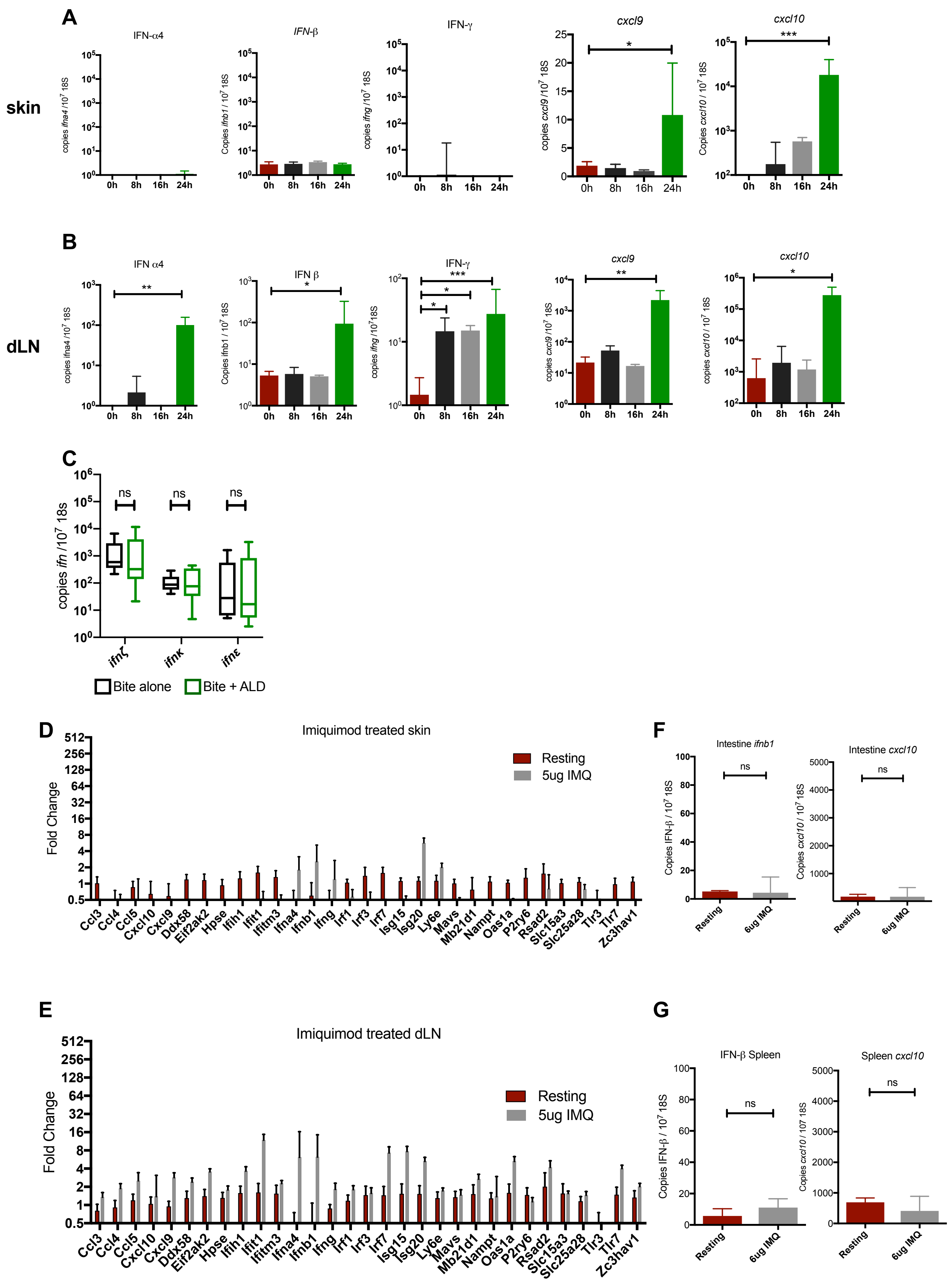
A
B
C
D
E
F
G
skin
dLN
IFN-α4
IFN-β
IFN-γ
cxcl9
cxcl10
IFN α4
IFN β
0h
8h
16h
24h
ns
ifnζ
ifnκ
ifnε
Bite alone
Bite + ALD
Imiquimod treated skin
Imiquimod treated dLN
Resting
5ug IMQ
Fold Change
Intestine ifnb1
Intestine cxcl10
IFN-β Spleen
Spleen cxcl10
Resting
6ug IMQ
Ccl3
Ccl4
Ccl5
Cxcl10
Cxcl9
Ddx58
Eif2ak2
Hpse
Ifih1
Ifit1
Ifitm3
Ifna4
Ifnb1
Ifng
Irf1
Irf3
Irf7
Isg15
Isg20
Ly6e
Mavs
Mb21d1
Nampt
Oas1a
P2ry6
Rsad2
Slc15a3
Slc25a28
Tlr3
Tlr7
Zc3hav1

**Fig. S2. Type I IFN and ISG expression in skin and draining LN following IMQ application.** (A-C). Mice were treated with 2 mg of topical ALD (5% IMQ) in the presence of a mosquito bite from 1-hour post bite. Copy number of host *ifna4*, *ifnb1*, *ifng*, *cxcl9*, *cxcl10* and 18S RNA transcripts in the skin (A), and draining popliteal lymph node (B), were determined by qPCR at 0h, 8h, 16h and 24h post ALD application (n=6). (C) Expression of atypical type I IFN were also assessed at 24 hours post ALD in the skin.

(D-G) Mice were injected with 6 µg of IMQ s.c. (D) Fold change in gene expression of ISG and IFN in skin (D) and dLN (E) was determined by qPCR at 24h post treatment (n=4). (F,G) Copy number of host *ifnb1*, *cxcl10* and 18S in the small intestine (F) and spleen (G) was determined by qPCR at 24h post treatment (n=6).

*p<0.05, **p<0.01, ***p<0.001, ns=not significant (Kruskal-Wallis test)

### Fig. S3

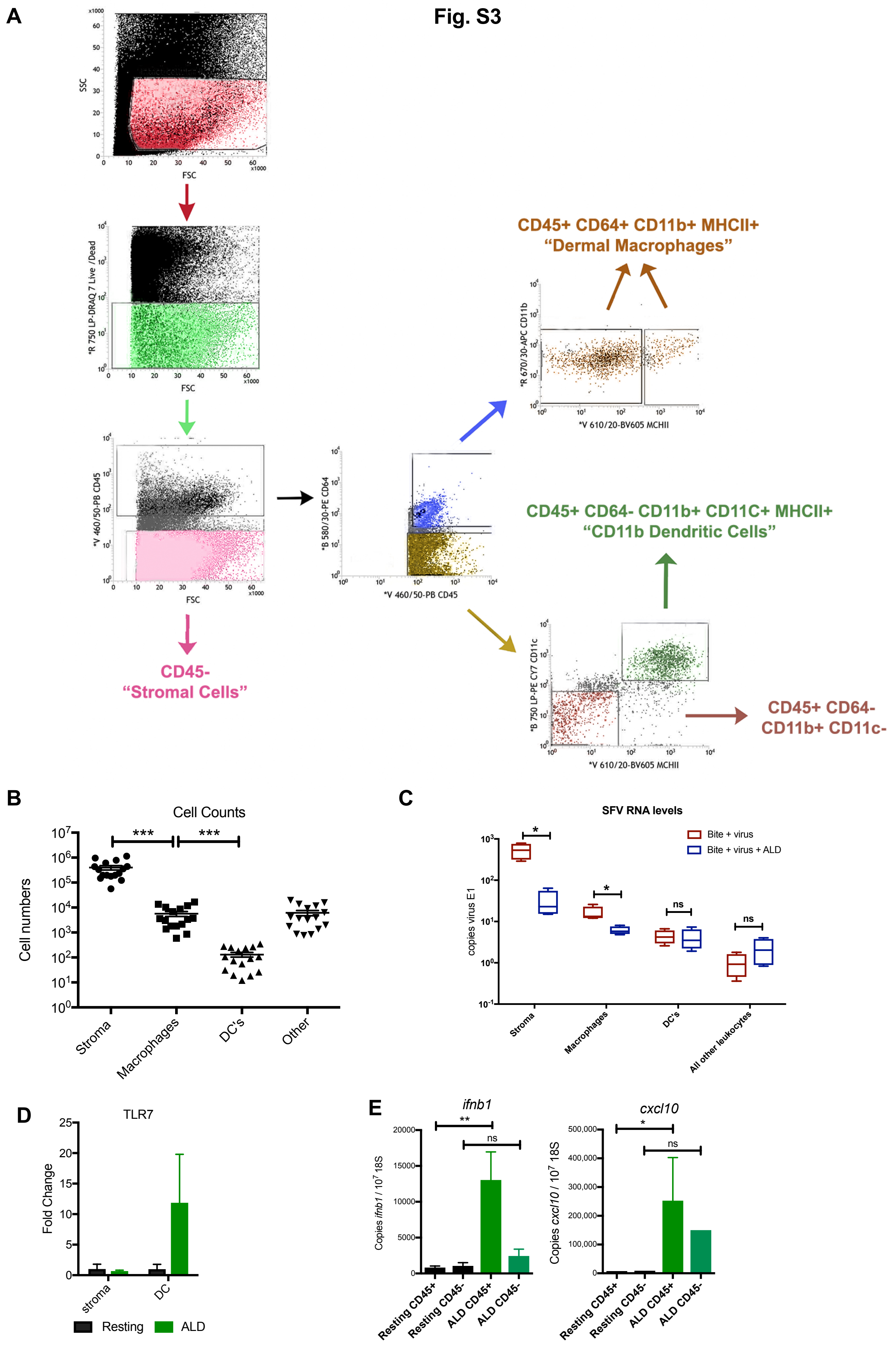

A
SSC
FSC
*R 750 LP-DRAQ 7 Live /Dead
*V 460/50-PB CD45
*B 580/30-PE CD64
*V 460/50-PB CD45
*R 670/30-APC CD11b
*V 610/20-BV605 MCHII
*B 750 LP-PE CY7 CD11c
*V 610/20-BV605 MCHII
CD45+ CD64+ CD11b+ MHCII+
“Dermal Macrophages”
CD45+ CD64- CD11b+ CD11C+ MHCII+
“CD11b Dendritic Cells”
CD45-
“Stromal Cells”
CD45+ CD64-
CD11b+ CD11c-
B
Cell Counts
Cell numbers
***
***
Stroma
Macrophages
DC's
Other
C
SFV RNA levels
Bite + virus
Bite + virus + ALD
copies virus E1
*
*
ns
ns
Stroma
Macrophages
DC's
All other leukocytes
D
TLR7
Fold Change
stroma
DC
Resting
ALD
E
ifnb1
Copies ifnb1 / 10^7 18S
**
ns
Resting CD45+
Resting CD45-
ALD CD45+
ALD CD45-
cxcl10
Copies cxcl10 / 10^7 18S
*
ns
Resting CD45+
Resting CD45-
ALD CD45+
ALD CD45-

**Fig. S3. Gene expression analysis of skin inoculation site-derived FACS isolated cells.**

(A-B) Gating strategy of flow cytometry sort. Mice were mosquito bitten and infected with SFV6. At 24 hpi, mouse skin was digested enzymatically in presence of transcriptional inhibitors, cells stained for lineage markers and FACS sorted at 4°C. Cells sorted into "stroma" (CD45-ve), "macrophage" (CD45+ve CD64+ve CD11b+ve, MHC$^{\text{low to high}}$), dermal dendritic cell "DC's" (CD45+ve CD64-ve CD11c$^{hi}$ CD11b+ MHC-II$^{hi}$) and "all other leukocyte" compartments, which are the bulk combination of all other remaining CD45+ve cells. Sorted macrophages were also positive for F4/80.

(B) Representative cell counts of a FACS sort. Stromal cells represented the most numerous cell type, while the macrophage population was significantly more numerous than DC population.

(C,D) Mice were infected with SFV6 s.c. in the presence of a mosquito bite, then treated with topical ALD from 1h post infection. At 24 hpi, skin was digested and cells FACS isolated. Copy number of SFV RNA (C) and *tlr7* (D) was determined by qPCR (n=4).

(E) Mice were bitten with mosquitoes and 1 hour later treated with topical ALD. At 24h post treatment skin was digested and cells sorted into "stroma" (CD45-ve) and "leukocytes" (CD45+ve). Copy number of host *ifnb1*, *cxcl10* and 18S were determined by qPCR (n=3). *$p<0.05$, **$p<0.01$, ***$p<0.001$, ns=not significant (Mann-Whitney test).

**Fig. S4**

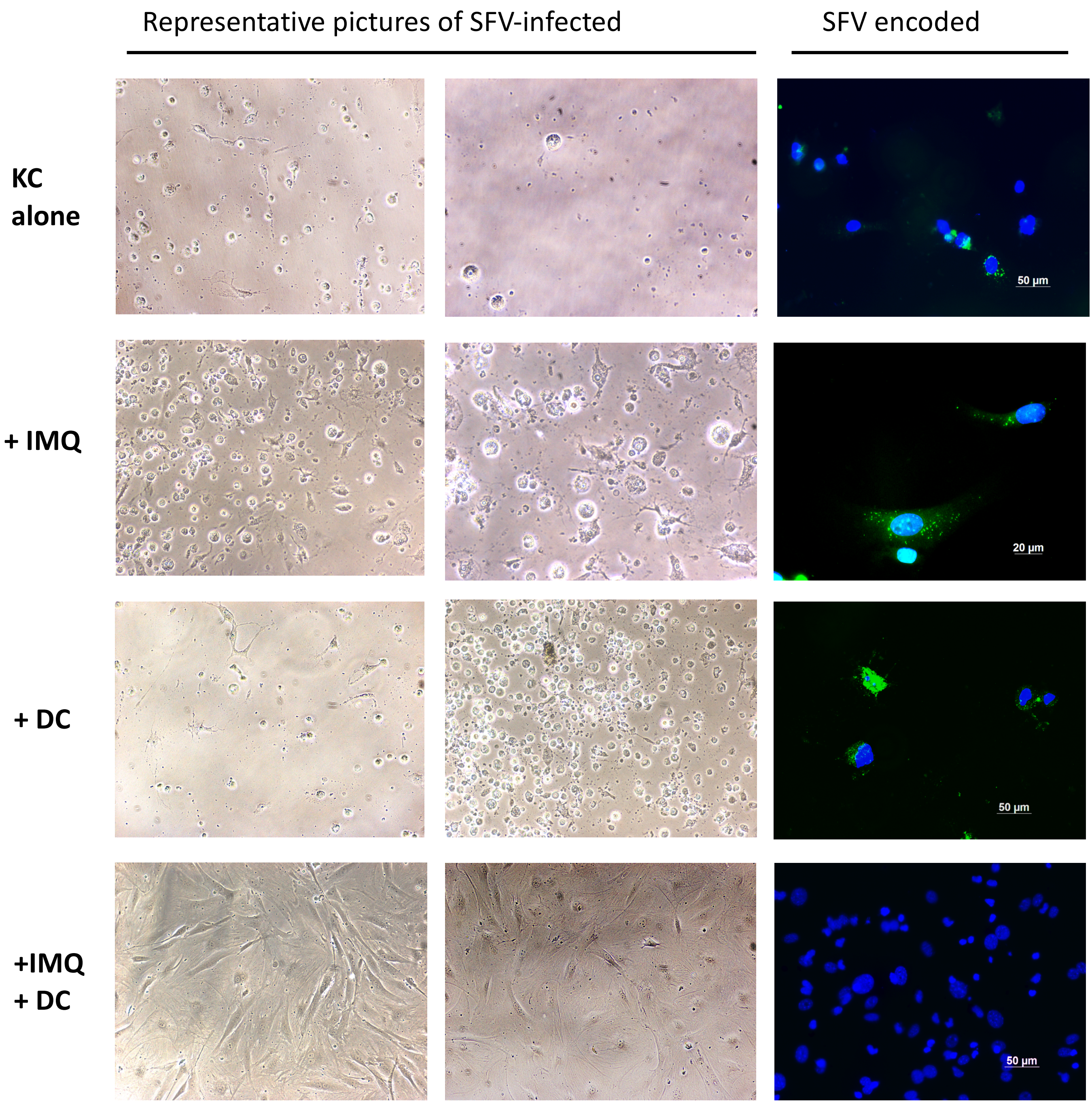


**Figure S4. IMQ-mediated protection against virus infection in keratinocytes is dependent on help from leukocytes.**

Monolayers of primary mouse keratinocytes were infected *in vitro* with SFV4(Xho)-EGFP (green) at an MOI of 0.1 in the presence or absence of DCs, then treated 1 hour later with 0.2 µg/ml IMQ. Cells were stained for dapi (blue).

Shown are representative images taken by brightfield and epifluorescence microscopy, to demonstrate extent of keratinocyte monolayer that remained intact at 24 hpi. In the absence of DC, monolayer was destroyed by virus-induced cell lysis. The few remaining keratinocytes that survived by 24hphi were positive for SFV-encoded eGFP. In the presence of DC, IMQ was able to preserve the keratinocyte monolayer, and little eGFP was detected.
